# TRPtracker: a community database for monitoring praziquantel sensitivity at TRPM_PZQ_ variants

**DOI:** 10.1101/2025.08.27.671753

**Authors:** Claudia M. Rohr, Sang-Kyu Park, Kelsilandia Aguiar-Martins, Timothy J.C. Anderson, Duncan J. Berger, Matthew Berriman, Sarah K. Buddenborg, Amaya L. Bustinduy, Frédéric D. Chevalier, James A. Cotton, Thomas Crellen, Stephen R. Doyle, Aidan M. Emery, Julien Kincaid Smith, Safari Kinung’hi, Poppy H. L. Lamberton, Winka Le Clec’h, Eric Ndombi, Tom Pennance, Candia Rowel, Shannan S. Summers, John Vianney Tushabe, Martin Walker, Bonnie L. Webster, Joanne P. Webster, Shona Wilson, Jonathan S. Marchant

## Abstract

The anthelmintic praziquantel (PZQ) has been used for decades as the clinical therapy for schistosomiasis, and remains the only available drug. As a cheap and effective drug therapy for all human disease-causing *Schistosoma* species, usage of PZQ underpins mass drug administration strategies aimed at eliminating schistosomiasis as a public health problem by 2030. Concern over the potential emergence of resistance to PZQ is therefore warranted, as it would constitute a major threat to this approach. In terms of molecular adaptations conferring PZQ resistance, variation in the sequence and/or expression of the drug target is an obvious mechanism and should be a priority for surveillance efforts. The target of PZQ is a transient receptor potential ion channel, TRPM_PZQ_, which is established as a locus that regulates schistosome sensitivity to PZQ. Here, we describe the establishment of a community resource, ‘TRPtracker’, which coalesces data on TRPM_PZQ_ natural variants together with measurements of individual variant sensitivity to PZQ. A compendium of laboratory-generated mutants in TRPM_PZQ_ is also compiled in TRPtracker to map regions within TRPM_PZQ_ critical for PZQ sensitivity. Aggregation of data from multiple research groups into TRPtracker permits rapid community-wide exchange of data, cataloguing which TRPM_PZQ_ variants have been functionally profiled, where geographically these variants have been found, their frequency within populations and their potential impact on PZQ sensitivity.

## Introduction

Schistosomiasis is a disease caused by infection with parasitic blood flukes of the genus *Schistosoma* [1]. Over 240 million people are estimated to be infected worldwide, and the current World Health Organization roadmap advances the goal of elimination of schistosomiasis as a public health problem, with interruption of transmission in selected regions, by 2030 [2, 3].

A key part of this strategy centers on mass drug administration (MDA) campaigns utilizing praziquantel (PZQ), an anti-schistosomal drug originally discovered in the 1970s. While an ‘old’ drug, PZQ has proven an effective therapy, both in terms of treatment costs and outcomes [4, 5]. However clinical dependence on a single therapy has nurtured lingering concern over the emergence of drug resistance. This concern has heightened following increased rollout of MDA centered on PZQ, with reduced drug efficacies already reported within human populations under the longest and strongest PZQ pressures [6]. This is further exacerbated through scenarios of poor compliance with dosing, inappropriate dosing [7], as well as non-ideal PZQ usage (for example treating livestock [2]). Given that decreased sensitivity to PZQ is a selectable trait in laboratory schistosome populations and schistosomiasis treatment failures have been reported in the literature, caution over whether reliance on PZQ will be sufficient to reach elimination goals appears prudent [8, 9]. Recent reports of PZQ resistance in cestodes from studies in companion animals and horses heightens concern over the potential for emergence of resistance [10–13].

One challenge in assessing and monitoring PZQ efficacy, has been a paucity of genetic markers associated with decreased PZQ sensitivity [14]. One reason for this is that the basis of PZQ action has long remained mysterious as no parasite target for PZQ was known. This picture has thankfully changed, following the recent discovery of an ion channel from the transient receptor potential melastatin subfamily (TRPM_PZQ_) that is activated by PZQ [15, 16]. Various lines of evidence support TRPM_PZQ_ as the clinically relevant target of PZQ in schistosomes and other parasitic flatworms sensitive to PZQ [17]. Identification of TRPM_PZQ_ as the parasite PZQ binding target defines an obvious locus for profiling PZQ sensitivity of natural TRPM_PZQ_ variants, consistent with a plethora of examples of ‘on-target’ mutations that confer decreased sensitivity to a variety of drugs [18, 19]. Indeed, leveraging variation that has occurred over an evolutionary timescale, it has been shown that the identity of a single amino acid within the PZQ binding pocket of *Fasciola spp*. TRPM_PZQ_ renders *Fasciola spp*. TRPM_PZQ_ refractory to PZQ [20, 21]. This is consistent with the insensitivity of these liver fluke to PZQ and the inability to cure fascioliasis with PZQ. Similarly, another instance of sequence variation within the TRPM_PZQ_ binding pocket of pseudophyllidean cestodes decreases sensitivity to PZQ into the micromolar range [21, 22], an observation consistent with the poor resolution of clinical infections caused by these tapeworms with standard PZQ dosing. Therefore, sequence variation within TRPM_PZQ_ dictates clinical treatment strategies for different types of infection.

These examples underscore the importance of sampling variation at the *TRPM_PZQ_* locus within natural schistosome populations, as variation could impact the efficacy of PZQ in treating schistosomiasis. Indeed, *Smp_246790*, the *Schistosoma mansoni* gene encoding TRPM_PZQ_ has been linked to decreased PZQ sensitivity in a laboratory population of *S. mansoni* selected for PZQ resistance [23, 24]. Clinically, we need to know whether poor schistosomiasis cure rates after multiple rounds of PZQ chemotherapy, often referred to as localized ‘hot spots’ of disease persistence, relate to the presence of specific TRPM_PZQ_ variants, and/or lowered expression of TRPM_PZQ_ within different schistosome populations. How extensive is standing genetic variation in TRPM_PZQ_? Do specific single nucleotide polymorphisms (SNPs), indels, copy number variants or more complex adaptations contribute to lower TRPM_PZQ_ responsivity to PZQ, or is natural variation in TRPM_PZQ_ completely benign without any impact on PZQ efficacy? Will comparable resistant mechanisms exist more broadly across the *Schistosoma* genus, encompassing other major disease-causing species such as *S. japonicum* and *S. haematobium*? The identification of TRPM_PZQ_ now facilitates all such analyses. This knowledge is critically important for design of effective schistosomiasis control and elimination strategies, enabling surveillance of the geographic distribution of any TRPM_PZQ_ variants of concern and informing PZQ treatment approaches in areas where such low sensitivity variants persist. Given the availability, low cost, and general effectiveness of PZQ as the sole treatment for schistosomiasis, any decline in the clinical effectiveness of this drug would prove a serious global health challenge.

Following identification of TRPM_PZQ_, it is now timely to profile the PZQ sensitivity of natural TRPM_PZQ_ genetic variants and ensure these data are broadly available to help surveillance efforts of PZQ effectiveness worldwide. In this manuscript, we report the establishment of a community resource (www.TRPtracker.live, [25]) which compiles TRPM_PZQ_ variant data contributed from multiple research groups in parallel with measurements of PZQ sensitivity and TRPM_PZQ_ expression. Combination of these data into a single resource permits easy reference of which TRPM_PZQ_ variants have been functionally profiled to date, where these variants have been found geographically, and whether any specific variants are associated with decreases in PZQ sensitivity.

## Methods

### Materials & Reagents

(±)-PZQ was purchased from Sigma (St. Louis, MO). All cell culture reagents were from Invitrogen (Waltham, MA). HEK293 cells (CRL-1573) were sourced from ATCC (Manassas, VA) and were tested monthly for mycoplasma contamination using the LookOut^®^ Mycoplasma PCR Detection Kit (Sigma).

### Functional assays of TRPM_PZQ_ variants

Coding sequence variants of TRPM_PZQ_ were made within the canonical reference TRPM_PZQ_ sequence (*Smp_246790.1*, WormBase Parasite, SM_V10, [26]) by Genscript (Piscataway, NJ) and confirmed by sequencing. TRPM_PZQ_ variant sensitivity to racemic PZQ ((±)-PZQ) was assessed using a cytoplasmic Ca^2+^ reporter assay using a high affinity Ca^2+^ dye (Fluo-4 NW, Thermo Fisher Scientific). This Ca^2+^ reporter assay was performed in black-walled, clear-bottomed 384-well plates previously coated with poly-L-lysine (Greiner Bio-One, Germany). Non-transfected, or TRPM_PZQ_ variant transfected, cells were seeded (20,000 cells/well) in individual wells and incubated in DMEM growth media containing 10% FBS. After 24 hours, this medium was replaced with 20µl of Fluo-4 NW dye loading solution (Molecular Devices) previously reconstituted in assay buffer (Hanks’ balanced salt solution containing 0.126 mM Ca^2+^, 0.49 mM Mg^2+^, 20 mM HEPES and 2.5 mM probenecid). Cells were incubated for 30 min at 37°C (5% CO_2_) followed by an additional 30 min incubation at room temperature. Changes in fluorescence were then resolved in real-time using a Fluorescence Imaging Plate Reader (FLIPR^TETRA^, Molecular Devices). Basal fluorescence (filter settings λ_ex_=470-495nm, λ_em_=515-575nm) from each well was captured for 20s, then (±)-PZQ (5μl) or vehicle solution (5μl), was added (25μl total volume) and the signal recorded over a subsequent 250s. Changes in fluorescence were represented as relative fluorescence units after subtracting the average basal fluorescence (averaged over 20s) from the recorded values. Concentration-response analysis was performed using four parameter sigmoidal curve fitting functions in Prism (v10.4.2, GraphPad Software, Boston, Massachusetts USA, www.graphpad.com)) from n≥3 technical replicates per mutant/variant. For constructs where data diverged from wild type *Sm*.TRPM_PZQ_ sensitivity, data collection was repeated from n≥3 independent transfections. Sensitivity of *Sm*.TRPM_PZQ_ to PZQ was profiled in response to a PZQ concentration range spanning 0.1nM to 100µM. The relative activity (RA) value describing PZQ sensitivity at individual TRPM_PZQ_ variants was calculated from the reference (wild type) vs variant half maximal effective concentration (EC_50_) and peak response (B_max_) using the following equation: RA = [EC_50,wild type_ x B_max,variant_] / [EC_50,variant_ x B_max,wild type_].

### Western Blotting

Cell lysates for Western blotting were prepared from HEK293 cells transfected with different TRPM_PZQ_ constructs. In brief, confluent cells were washed twice with ice-cold PBS, harvested into 1 ml of PBS, and pelleted at 1,000 x g for 5 minutes. The cell pellet was then resuspended in 300-500μl of lysis buffer (1x PBS, 1% Triton) containing a complete protease inhibitor mixture (Roche Applied Science). Samples were then incubated for 30 minutes at 4℃ on a rotator. The sample was then centrifuged at 10,000 x g for 10 minutes, and the supernatant used for Western blot analysis. Protein concentrations were determined using a BCA assay (Bio-Rad) and equal amounts of protein (20–30μg) were loaded into individual lanes of a 4-15% Tris-glycine gel (Bio-Rad). After electrophoresis, samples were transferred to a PVDF membrane, blocked with blocking buffer in (5% BSA, 0.1% Tween/Tris buffered saline (TBST)) for 1 hour at room temperature. The membrane was incubated overnight at 4°C with either a custom TRPM_PZQ_ antibody (1:1000) raised against the COOH-terminal region of TRPM_PZQ_ (1709-1997 amino acids), or a β-actin antibody (1:2000). After washing with TBST, the membrane was incubated (1 hour, room temperature) with a horseradish peroxidase-conjugated secondary antibody (Invitrogen). Following additional washes, protein bands were visualized using a chemiluminescent substrate (ECL, Invitrogen) and quantified using the ChemiDoc™ MP Imaging System (BioRad).

## Results

The TRPtracker database [25] collates data reporting the PZQ sensitivity of individual TRPM_PZQ_ variants. These data have been compiled from two sources (**Figure 1A**). The first dataset represents a collection of ‘laboratory’ TRPM_PZQ_ mutants made to probe the structure-function relationship of TRPM_PZQ_ [15, 20]. The second dataset represents natural TRPM_PZQ_ variants, informed by sequencing data from natural schistosome populations that have been prioritized for functional evaluation by various research groups. Both datasets provide utility to this resource. While lab generated mutants impart no clinical relevance, they do inform about regions of TRPM_PZQ_ relevant for function which may in turn guide selection of natural variants for functional profiling from large sequencing datasets. At the time of writing, both datasets establish a ‘library’ of ∼170 TRPtracker database entries. Each entry is clearly labelled as to their derivation (i.e. lab mutant (‘lab’) or natural variant (‘field’)). Database entries have been limited to those variants that have been subjected to functional profiling.

**Figure 1.**
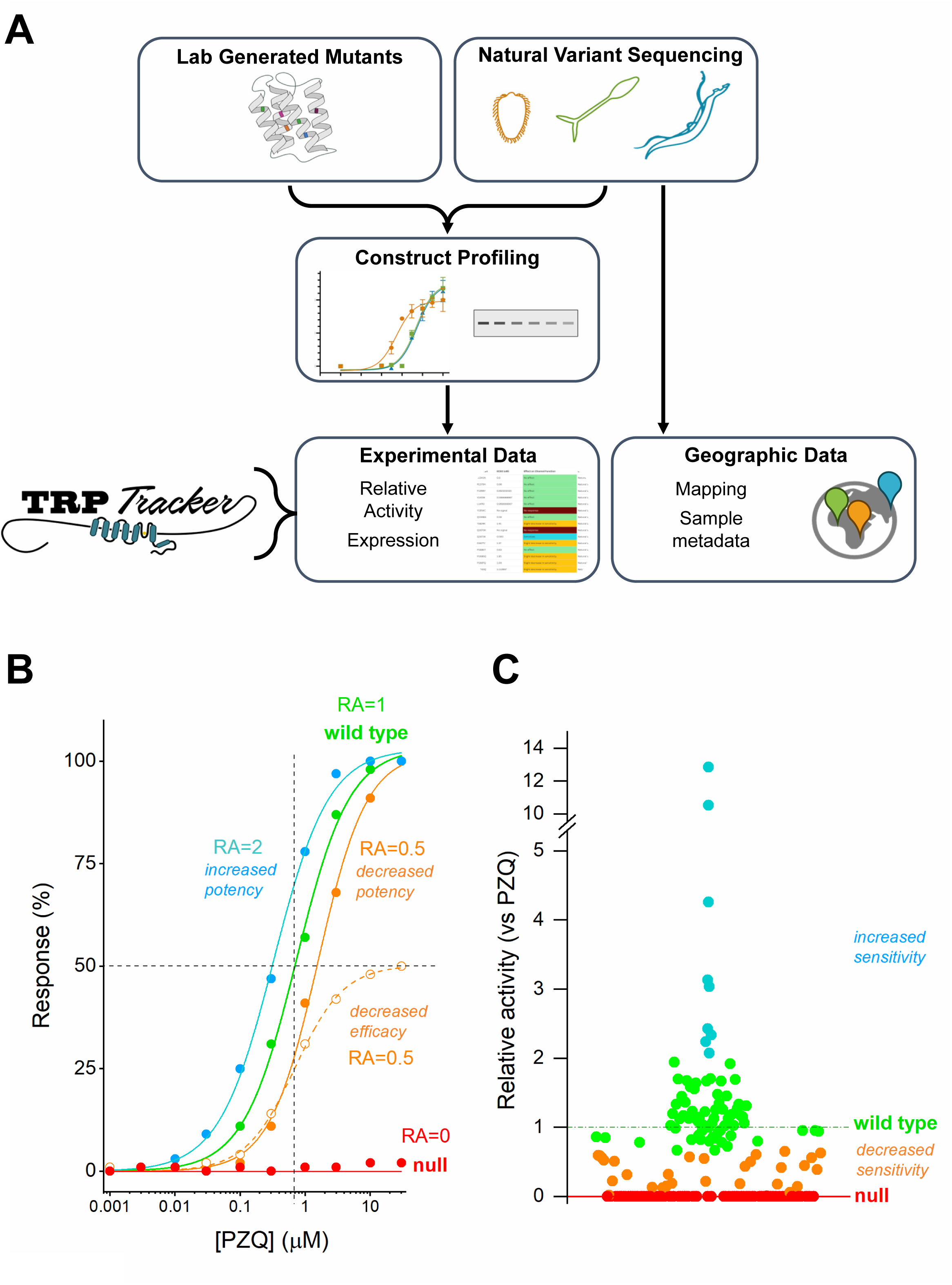
Schematic overview of TRPtracker and functional profiling of TRPM_PZQ_. (**A**) Schematic overview of profiling workflow for lab-generated mutants and natural variants, which can be profiled functionally and in terms of expression levels. These data and sample metadata are compiled in the TRPtracker resource. (**B**) Example concentration response curves to illustrate impact of different TRPM_PZQ_ variants on the relative activity (RA) of PZQ. Increases in potency yield higher RA values (blue), decreases in potency or decreases in efficacy yield lower RA values (orange) relative to the wild type TRPM_PZQ_ concentration response curve (green). (**C**) Relative activity plot of *Sm*.TRPM_PZQ_ variants from the TRPtracker dataset, stratified into different function categories: null (RA = 0, red), decreased sensitivity (RA = 0.6 -1, orange), wild type (RA = 0.6 -1.4, green) and increased sensitivity (RA> 1.4, blue).

Functional profiling was performed using an *in vitro* Ca^2+^ reporter assay based upon heterologous expression of individual TRPM_PZQ_ variants in HEK293 cells. TRPM_PZQ_ is a non-selective cation channel, and changes in Ca^2+^ concentration resulting from TRPM_PZQ_ activity can be resolved using a fluorescent Ca^2+^ indicator dye loaded into cells. The amplitude of PZQ-evoked Ca^2+^ signals correlate with the sensitivity of TRPM_PZQ_ activation by PZQ, which displays an EC_50_ in the sub-micromolar range for the reference *Schistosoma mansoni* TRPM_PZQ_ sequence (EC_50_ of 739±171nM for (±)-PZQ at *Sm*.TRPM_PZQ_). A Western blotting assay to resolve TRPM_PZQ_ protein expression has also been used for a subset of variants to probe the basis of changed PZQ sensitivity. Experimental data have been compiled together with sample metadata, which for natural variants typically includes information about the sample collection, including geographical foci and any recent PZQ treatment history. This profiling workflow is summarized schematically in Figure 1A.

The functional impact of individual mutants is quantified as a ‘relative activity’ measure (RA, [27]) relative to PZQ action at the *Sm*.TRPM_PZQ_ reference sequence. The RA parameter was chosen as it incorporates how each mutant impacts PZQ potency (left/right shift in curve EC_50_) as well as PZQ efficacy (up/down shift in curve maximum, B_max_) relative to values measured at the reference ‘wild type’ TRPM_PZQ_. This is important as changes in either parameter – ligand potency or efficacy – will impact overall worm sensitivity to PZQ. Mutants that worsen potency or efficacy result in RA values <1, whereas mutants that sensitize TRPM_PZQ_ to PZQ result in RA values >1. For example, a sensitizing mutant that increases PZQ potency by 2-fold equates to a RA value of 2. In contrast, a mutant that decreases the measured EC_50_ of PZQ at TRPM_PZQ_ by 2-fold or a mutant that decreases the efficacy of PZQ by 50% equate to a RA value of 0.5 (**Figure 1B**). The distribution of RA values for the current compendium of mutants is shown in **Figure 1C**. The measured RA values span a broad range, with a mean of ∼1. Data were stratified into functional categories with the distinction between ‘wild type’ and ‘decreased PZQ sensitivity’ groups based on the standard deviation for the population of natural variants. These categories comprised: ‘null’ (RA = 0, red), ‘decreased PZQ sensitivity’ (RA >0 but <0.6, orange), ‘wild type’ (RA, 0.6 – 2, green) and ‘sensitized’ (RA, >2, blue). All data, including sample metadata, are available for reference at the portal (www.TRPtracker.live, [25]). Functional profiling data are summarized in Table 1 and can also be downloaded at the TRPtracker website [25]. Insight from the ‘lab’ mutants and ‘field’ variants that have been functionally profiled are discussed in the following two sections.

### TRPM_PZQ_ laboratory mutants

Ongoing functional analyses of TRPM_PZQ_ has generated a collection of ∼140 point mutants and truncations within the reference sequence of *Sm*.TRPM_PZQ_. The distribution of these point mutants is currently biased to the transmembrane spanning regions of TPRM_PZQ_ (∼60% of mutants in the voltage sensor-like domains (VSLD), pore and TRP domains) because of prior work to define the binding sites of the TRPM_PZQ_ ligands PZQ and BZQ [20, 28]. Multiple ‘null’ mutants that completely ablated PZQ sensitivity were identified (**Figure 2A**), many of which were localized within the transmembrane spanning domains of TRPM_PZQ_. While this weighting reflects the aforementioned bias in studying this region of TRPM_PZQ_, the data do underscore the criticality of the VSLD architecture (S1-S4, containing the PZQ binding pocket) pore domain structure (S5-S6), the TRP helix and their intervening connections for normal channel function.

**Figure 2.**
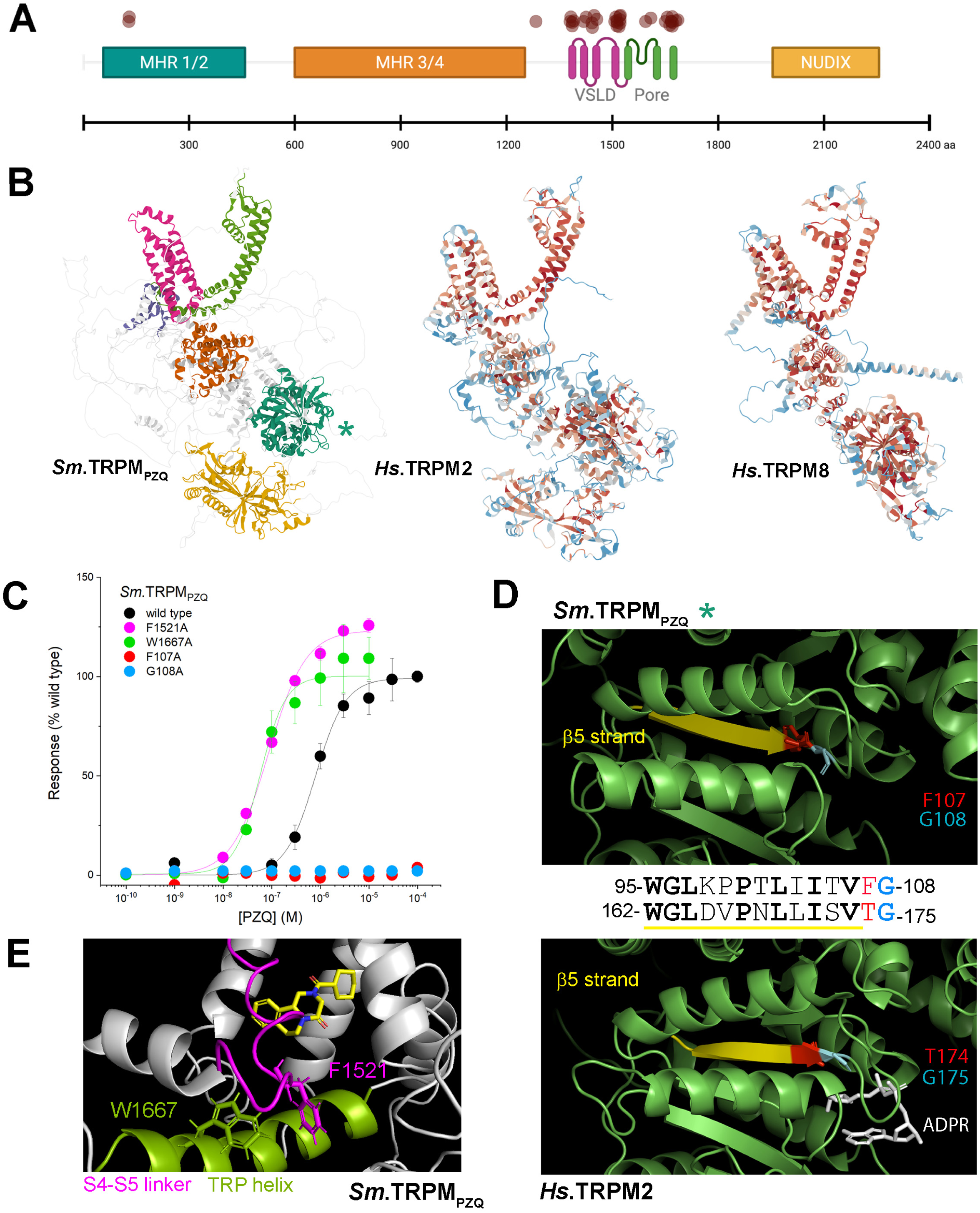
Functional effects of various TRPM_PZQ_ mutants. (**A**) Distribution of functionally ‘null’ laboratory mutants (red circles) along the length of the *Sm*.TRPM_PZQ_ coding sequence. (**B**) Representations from the AlphaFold protein structure database [56, 57] for a monomer of *Sm*.TRPM_PZQ_ (left), *Hs*.TRPM2 (O94759, middle) and *Hs*.TRPM8 (Q7Z2W7,right). The *Sm*.TRPM_PZQ_ prediction shows several predicted structural domains [58] corresponding to the voltage-sensing like domain (VSLD, S1-S4, magenta), pore domains (S5-S6, green), MHR3/4 (orange), MHR1/2 (turquoise), nudix hydrolase domain (yellow). Overall, *Sm*.TRPM_PZQ_ presents with homology to the vertebrate TRPM8-like menthol binding pocket found in the VSLD (S1-S4) within a broader protein structure with cytoplasmic domain homology to *Hs*.TRPM2 [16]. The monomers of *Hs*.TRPM2 and *Hs*.TRPM8 are colored in terms of predicted effect on protein ‘pathogenicity’ function using AlphaMissense [29], with benign variation colored blue and deleterious mutants scored with increasing warm coloration toward red. (**C**) Functional profiling of *Sm*.TRPM_PZQ_ mutants in response to PZQ. These include null mutants (*Sm*.TRPM_PZQ_ [F107A] and *Sm*.TRPM_PZQ_[G108A]) as well sensitizing mutants (*Sm*.TRPM_PZQ_ [F1521A] and *Sm*.TRPM_PZQ_[W1667A]). (**D**) Interrogation of a null mutant ‘hot-spot’ in the NH_2_-terminal MHR1/2 domain of TRPM_PZQ_. The null mutants found in *Sm*.TRPM_PZQ_ profiled in (C) localize to the end of a β-strand (F107 in red, G108 in cyan, top) which shows structural and sequence conservation with the β5-strand in *Hs*.TRPM2 (T174 in red, G175 in cyan; bottom, [59]). This region of TRPM2 is implicated in the binding of ADP-ribose [59], shown in white (PDB, 8E6V). (**E**) Projection of location of the sensitizing mutants *Sm*.TRPM_PZQ_ [F1521A] (magenta, S4-S5 linker) and *Sm*.TRPM_PZQ_[W1667A] (green, TRP domain) relative to the PZQ (yellow) in the VSLD binding pocket.

However, it is important to recognize that mutations in other regions of TRPM_PZQ_ will also compromise PZQ action. Considering examples of human TRPM channels, the distribution of *in silico* predicted deleterious mutants spans multiple protein domains throughout the entire coding sequence. Predictions using the ‘pathogenicity’ function of AlphaMissense [29] are shown in **Figure 2B** for human TRPM2 (*Hs*.TRPM2) and human TRPM8 (*Hs*.TRPM8), both of which display structural conservation with TRPM_PZQ_ [16, 20]. In the human TRPM models, benign variation is colored blue and deleterious mutants are scored with increasing warm coloration. As evident through this structural comparison, predicted deleterious mutants span multiple domains within the human TRPM channels beyond the transmembrane spanning region, throughout domains that show conservation with TRPM_PZQ_ (Figure 2B).

Consistent with this prediction, a cluster of functionally ‘null’ mutants was evident at the NH_2_-terminus of *Sm*.TRPM_PZQ_ within the experimental dataset. For example, the dual mutants *Sm*.TRPM_PZQ_[F107A] and *Sm*.TRPM_PZQ_[G108A] exhibited no responsiveness to PZQ (**Figure 2C**). These residues are found in a region of the MHR1/2 domain that is well conserved between TRPM_PZQ_ orthologs (Supplementary Figure 1). Structural comparison of this region in *Sm*.TRPM_PZQ_ compared with *Hs*.TRPM2 shows these dual residues are located at the end of a β-strand (β5 in the *Hs*.TRPM2 MHR1/2 domain) which displays a similar predicted folding structure to the similar region of *Sm*.TRPM_PZQ_ (**Figure 2D**). Interestingly, the β5 strand in *Hs*.TRPM2 is known to form important interactions with an endogenous ligand, ADP ribose, which serves as an obligatory *Hs*.TRPM2 co-agonist while also conferring redox sensitivity to the ion channel [30, 31]. These data support further investigation of the role of endogenous nucleotides in regulating TRPM_PZQ_ expression and/or function [32]. This could be of interest in light of the customized nucleotide metabolic pathways present in schistosomes [33, 34].

The highest sensitizing mutants to PZQ in *Sm*.TRPM_PZQ_ (RA values of >10) Figure 1C) merit comment. These two mutants were *Sm*.TRPM_PZQ_[F1521A] (RA ∼10.5) and *Sm*.TRPM_PZQ_[W1667A] (RA ∼13, Figure 1C). Both mutants increased PZQ potency into the <100nM potency range. While each residue occurs in a different region of *Sm*.TRPM_PZQ_ (F1521 lies within the S4-S5 linker, W1667 is located in the TRP helix), they appear as potentially interacting residues (**Figure 2E**). This interaction likely serves as a ‘brake’ on ligand-evoked conformational change between the VSLD and pore domain that is necessary for channel opening. Consequently, ablation of either residue by alanine mutation serves to release this brake, more efficiently transducing ligand binding energy into the open state transition. The double mutant *Sm*.TRPM_PZQ_ [F1521A][W1667A] did not show enhanced potency to PZQ (EC_50_ of 68±11nM) consistent with the idea that these residues function in concert. Both these residues are well conserved in TRPM_PZQ_ orthologs (Supplementary Figure 2), suggesting this interplay may be a generalized feature of TRPM_PZQ_ activation.

In summary, these examples of ‘null’ and ‘sensitizing’ TRPM_PZQ_ mutants, highlight the principle that single residue variation found across the entire TRPM_PZQ_ coding sequence can change TRPM_PZQ_ sensitivity to PZQ.

### TRPM_PZQ_ field variants

Studies of laboratory mutants, while informative for understanding how *Sm*.TRPM_PZQ_ works, have no relevance to clinical infections unless similar variation is present and prevalent in the field. Importantly, several groups are currently analyzing TRPM_PZQ_ genetic data within natural schistosome populations to assess the scope of variation in this drug target [23, 35–37], and thereby relevance for clinical PZQ efficacy. Readers are referred to these individual studies for information regarding samples collected and the specific variants prioritized for functional profiling from sequencing analyses. Most of the genomic analyses have been performed fro*m S. mansoni* miracidia, isolated and preserved after hatching from eggs isolated from fecal samples.

From all of this ongoing work, ∼30 natural coding variants have now been profiled and entered into the TRPtracker database [25]. Many other variants have been reported but have not yet been functionally profiled. Data from the profiled variants, color coded in terms of sensitivity to PZQ is shown in **Figure 3A**, together with their geographical distribution in **Figure 3B**. Several of these variants have been found in samples collected and/or analysed by the different research groups. The vast majority of these variants (∼75%) exhibit, by our stratification of functional classes, no change in sensitivity toward PZQ (Figure 3A), with the average RA of the entire natural variation collection being 1.2±0.55 (mean±sd, 29 variants profiled to date). These are reassuring observations in the context of surveillance for ‘resistant’ variants within natural *S. mansoni* populations sampled to date.

**Figure 3.**
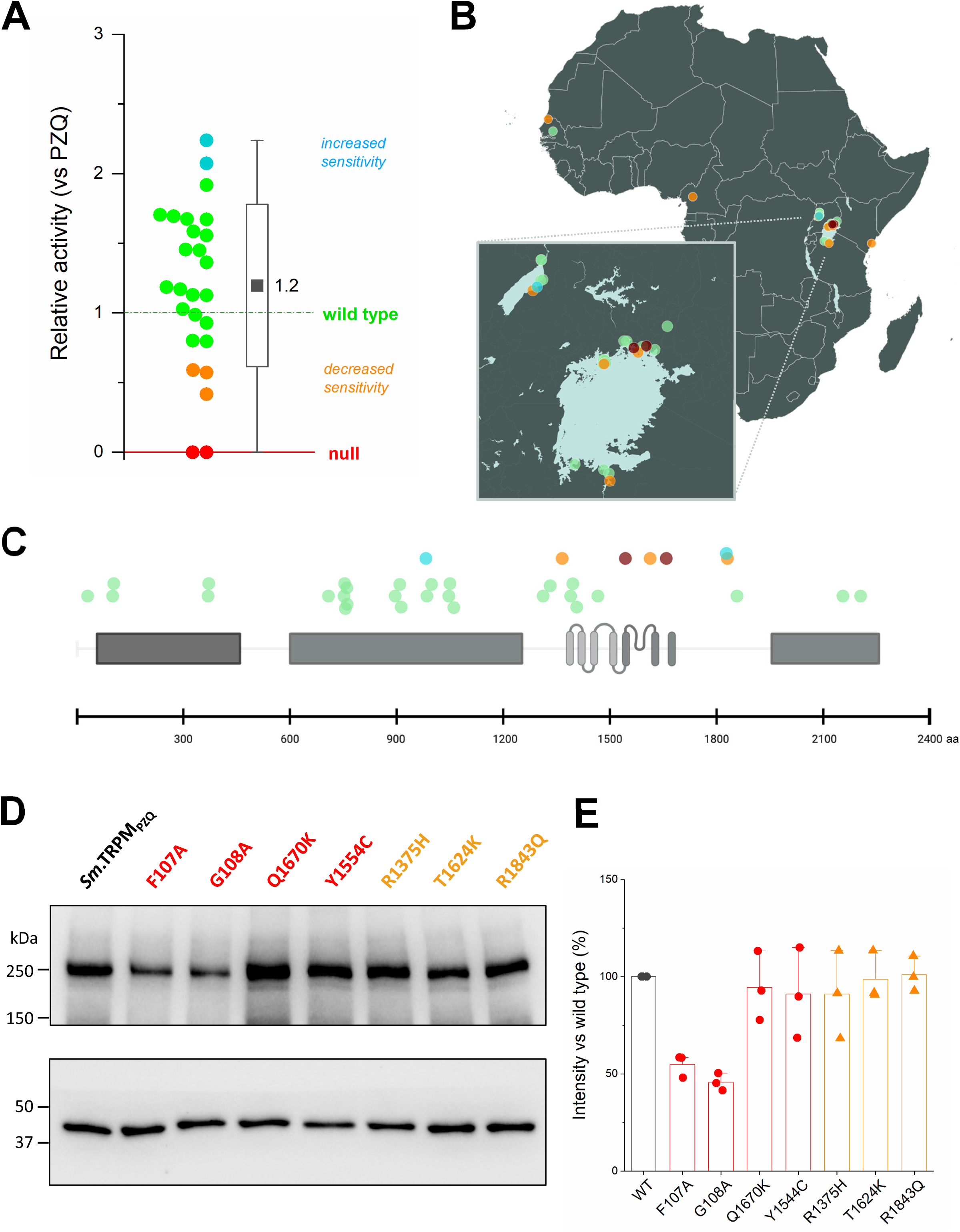
Profiling function and expression of various natural TRPM_PZQ_ variants. (**A**) Distribution of RA values for natural TRPM_PZQ_ variants that have been functionally profiled in in the database. Data is presented as a half-box plot, with data on the left and the standard deviation (box), range (vertical line) and mean (black square) of this population shown on the right. (**B**) Collection sites where the natural TRPM_PZQ_ variants that have been functionally profiled in this study were sampled. (**C**) Distribution of normal sensitivity (green), decreased sensitivity (orange) and functionally ‘null’ natural variants of *Sm*.TRPM_PZQ_ (red circles) identified by various groups along the length of the *Sm*.TRPM_PZQ_. (**D**) representative Western blot for expression of indicated TRPM_PZQ_ variants. (**E**) densitometric measurements from independent blots (n=3) quantified for these same TRPM_PZQ_ variants.

However, the dataset shown in Figure 3A does provide examples of natural variants which are functionally ‘null’ (Y1554C, Q1670K) or exhibit low RA values (<0.6, outside the standard deviation of the entire group, R1375H, T1624K, R1843Q). These lower sensitivity variants are mostly proximal (e.g. the pre-S1 helix) or within the transmembrane spanning region of TRPM_PZQ_ (**Figure 3C**). It is important to emphasize that many of these variants stratified into the ‘null’ or ‘impaired function’ categories are found at very low frequency in miracidial samples collected to date [23, 35–37]. Where data is available in the original reports, information about variant frequency is being incorporated into the TRPtracker resource. Unless a variant is identified in several samples at a specific site, or found by different groups across multiple sampling locales, caution is warranted as to whether these are *bona fide* natural variants or simply false calls from sequencing errors or amplification biases. Further sequencing and surveillance efforts will be of paramount importance to verify the occurrence and frequency of any predicted ‘variants of concern’ within natural worm populations.

Finally, another route toward manifestation of a lower worm sensitivity to PZQ is decreased expression of TRPM_PZQ_. Expression of all these natural TRPM_PZQ_ variants, as well as lab mutants *Sm*.TRPM_PZQ_[F107A] and *Sm*.TRPM_PZQ_[G108A], was therefore examined by Western blotting following transient transfection in HEK293 cells (**Figure 3D**). Expression levels were assessed by densitometry relative to the intensity of the ∼250kDa band in the *Sm*.TRPM_PZQ_ reference sequence (**Figure 3E**). Each natural variant that exhibited reduced PZQ sensitivity (R1375H, T1624K, R1843Q) or no PZQ sensitivity (Y1554C, Q1670K) displayed no difference in total TRPM_PZQ_ expression. These data suggest these variants impair *Sm*.TRPM_PZQ_ sensitivity to PZQ rather than *Sm*.TRPM_PZQ_ expression. This contrasts with the lack of function of the ‘null’ NH_2_-terminal lab mutants (*Sm*.TRPM_PZQ_[F107A] and *Sm*.TRPM_PZQ_[G108A], Figure 2C) where decreased *Sm*.TRPM_PZQ_ expression was evident by Western blotting (Figure 3D&E). This suggests a role for these NH_2_-terminal residues in regulating folding, stability or multimerization of TRPM_PZQ_. Overall, changes in either TRPM_PZQ_ expression or absolute sensitivity to PZQ underpin the lack of PZQ activity observed for the ‘null function’ TRPM_PZQ_ variants.

## Discussion

As a consortium of schistosomiasis researchers, we advance TRPtracker [25] as an online resource for collating data on the distribution and PZQ sensitivity of natural *TRPM_PZQ_* sequence variants. Our vision is that this database will be regularly updated with data from ongoing field collected *Schistosoma* samples and sequencing efforts being performed by several groups, such that data can be easily shared and accessed by the community. Data entries will include pertinent metadata, for example variant frequency at collection sites, sample descriptors, collection dates and PZQ treatment histories. Contributors are encouraged to submit such details through the website portal. Community input for additional suggestions to integrate into this resource are also welcome. The intent is that aggregating this information on a single platform should minimize redundancy by not reprofiling variants already characterized, while maximizing comparability across datasets by using standardized assays performed under identical conditions for all variants to allow side-by-side comparison of data. Currently, this database contains data for the PZQ sensitivity of ∼170 different *Schistosoma mansoni* TRPM_PZQ_ variants. However, it is envisaged that data entries will in time expand to include functional data on other *Schistosoma* species that infect humans and livestock worldwide, allelic variants of TRPM_PZQ_, and finally data addressing the impact of specific variants on TRPM_PZQ_ expression. These data could eventually be folded into an existing resource (such as the variant function of the Ensembl platform) or remain as a standalone catalogue like other initiatives that track drug resistance markers [38, 39].

TRPtracker is timely as sequencing efforts to define TRPM_PZQ_ variants are currently being performed by several groups. Readers are referred to these individual studies for interpretation of functional profiling data in the context of particular samples [23, 35–37]. Some speculative discussion is nevertheless possible as to what this ongoing work will likely reveal, by drawing upon existing knowledge of vertebrate TRP channel evolution as well as prior studies of polymorphism frequency within human TRP channels. We advance three predictions, justified in the following sections.

: first, considerable variation in TRPM_PZQ_ will be resolved and this variation will not be evenly spread across the gene,

: second, established precedent that TRP channel polymorphisms alter ligand sensitivity without impairing channel function portend a similar scenario for TRPM_PZQ_ unless fitness costs preclude selection of such variants,

: third, current analyses of coding variation within TRPM_PZQ_ need advance to anticipate the impact of non-coding variation, where potentially greater risk for PZQ ‘resistant’ phenotypes reside effected through changes in TRPM_PZQ_ expression levels.

### The landscape of TRPM_PZQ_ coding variants

TRPM_PZQ_ is a long TRP (2268 amino acids) encoded by a gene with 36 exons. A human TRPM that is of similar length is hTRPM6 (2022 amino acids, 39 exons) and over 72,000 variant alleles of hTRPM6 have been catalogued to date. TRPM_PZQ_ shares homology with both hTRPM8 (transmembrane binding pocket [20]) and hTRPM2 (COOH terminal NUDT9H homology [16], Figure 2B). Over 45,000 variant alleles have been described within both these human genes. In terms of coding sequence variation within hTRPM8, over 1,000 missense variations, at least 50 premature stop codons (the majority removing the entire transmembrane domain) as well as 70 frameshift mutations have been identified. This variation is distributed unevenly across the hTRPM8 coding sequence, with less variation evident within the transmembrane spanning region compared to the NH_2_ terminus [40, 41]. This constraint likely reflects the criticality of the voltage sensor-like domains (VSLD, transmembrane helices S1-S4), the pore domain (S5-S6) and the TRP helix/TRP box for channel function which exhibits higher constraint and therefore lower tolerance for variation. SNPs found in this region will have a higher probability of being deleterious to channel function. Similarly, from an evolutionary perspective, the transmembrane regions of TRPM8 are highly conserved through vertebrate evolution with very high levels of sequence conservation in TM4, TM5 and TM6 and the TRP box [41]. Such evolutionary scale analyses also implicate co-evolution of residues that form networks of allosteric regulation between the VSLD and pore domain of hTRPM8, again underscoring the importance of these regions to channel function [40].

More broadly across the coding sequence, conservation between different channel regions varies, reflective of different selection pressures during vertebrate TRPM8 evolution [40, 41]. This does not imply that regions with higher variability beyond the transmembrane spanning domain are benign (Figure 2B). Various diseases (TRP ‘channelopathies’ [42]) resulting from ‘gain-of’ and ‘loss-of’ -function mutants that perturb channel expression and/or regulation are found throughout the entire coding sequence of TRP channels. Functional profiling efforts for TRPM_PZQ_ have already identified a cluster of NH_2_-terminal mutants that ablate the PZQ sensitivity of TRPM_PZQ_ (Figure 2). In short, considerable variation will likely be evident and functional profiling across lab mutants, natural variants, and TRPM_PZQ_ splice variants [43] which help advance our understanding of how this ion channel works.

### Sequence variation and ligand sensitivity

In studies of vertebrate TRP channels, many examples have emerged of amino acid variation impacting sensitivity to environmental cues. For example, the temperature sensitivity of thermosensitive TRP channels has been correlated with sequence variation between different species [44–47]. The sensitivity of TRP orthologs in different species to natural products also varies between species. For example, the loss of capsaicin sensitivity in birds and a specific mammal (hTRPA1, [48]), or changes in menthol sensitivity during vertebrate evolution (hTRPM8, [46]). All such changes likely reflect selection of adaptive advantage as the sensory role of TRP channels attunes to the environmental milieu. That specific residues mediate sensitivity to discrete agonists is supported by a wealth of mutagenesis data, often guided by observation of species-specific insensitivities – for example icilin insensitivity in birds [49], and, of particular relevance to this topic, the PZQ insensitivity of *Fasciola* spp. TRPM_PZQ_ [20].

These studies demonstrate that TRP channels can lose sensitivity to specific chemotypes, without impairment of responsiveness to other activators or overall channel function. This underscores the polymodality of TRP channel activation where multiple modes of channel activation co-exist. This is true for TRPM_PZQ_, as the channel can be activated by different ligands as well as by membrane stretch [50]. Therefore, while it seems likely that PZQ-insensitive TRPM_PZQ_ variants will emerge in schistosome populations, the critical consideration is whether selection of such variants will confer a deleterious fitness cost preventing their expansion in natural populations [51]. TRPM_PZQ_ is known to depolarize neurons [50] and channel function may be essential for sensory and locomotory functions critical for life cycle progression.

Finally, thinking simply in terms of ‘gain’ or ‘loss’ of PZQ sensitivity is likely insufficient. Even small decreases in the PZQ sensitivity of schistosome TRPM_PZQ_ could reduce cure rates and be clinically significant. Quantitatively, it is the sensitivity of TRPM_PZQ_ to PZQ relative to PZQ exposure (the pharmacokinetic profile of PZQ concentrations in host mesenteric vessels after dosing) that is critical for the action of PZQ against adult worms. Even a slight decrease in TRPM_PZQ_ sensitivity to PZQ will shrink the effective kill zone within the PZQ pharmacokinetic profile that mediates worm elimination.

### Non-coding variation

The current focus on variation within the coding sequence of TRPM_PZQ_ will reveal only a small fraction of the variation associated with the *TRPM_PZQ_*gene. Non-coding variants, spanning the introns and UTRs of TRPM_PZQ_, as well as neighboring regulatory and structural variants will contribute the majority of variation and many of these variants may prove relevant in terms of PZQ sensitivity. Of note is the study of Le Clec’h *et al.* [23], who mapped markers associated with decreased PZQ sensitivity to regions close to, but not within *TRPM_PZQ_*, which were speculated to impact full length TRPM_PZQ_ expression. An example relevant for human TRP channel function is a non-coding SNP (rs10166942, C/T) found ∼1kb upstream of the human TRPM8 coding sequence This SNP displays one of the strongest genome-wide associations with migraine in humans [52, 53], with the ancestral ‘C’ allele being associated with reduced migraine risk and lower hTRPM8 expression [54, 55]. The derived ‘T’ allele is found at higher frequency in human populations in a latitude-dependent manner with higher TRPM8 expression conferring stronger, and presumably protective, cold temperature responsivity in cold-climates despite association with an increased migraine risk observed in European populations. This example underscores how selection of a non-coding variant has impacted TRP channel expression and activity across distinct populations. In a similar vein, studies of non-coding *TRPM_PZQ_* variants that impact channel expression will be important for understanding schistosome sensitivity to PZQ.

In conclusion, the TRPtracker resource establishes a community-driven portal to progressively compile data on the genetic variation within TRPM_PZQ_ across natural *Schistosoma* populations together with the functional impact of these variants. The resource will permit surveillance of whether drug target-related changes in PZQ sensitivity exist in natural schistosome populations and, if so, help monitoring the spread of specific variants of concern, allowing more tailored control interventions to be launched.

## Acknowledgements

Research was supported by the National Institutes of Health R01-AI145871 (to JSM), F31-AI183573 (to CMR), R01-AI123434 (to TJCA), R01-AI160433 (to EN). The FibroScHot project is part of the EDCTP2 programme supported by the European Union (RIA2017NIM-1842-FibroScHot). Collections associated with the Natural History Museum (NHM) in London, were supported by: EU grant CONTRAST (FP6 STREP contract no: 032203), NHM internal funds, and the Schistosomiasis Collection at the NHM (SCAN; Wellcome Trust biomedical resource project, 104958/Z/14/Z). The Schistosomiasis Consortium for Operational Research and Evaluation (SCORE) program was funded by the University of Georgia Research Foundation, Inc., which was funded by the Bill & Melinda Gates with the collections funded via sub-awards [RR374-053/5054146; RR374-053/4785426]. Other sample collections were supported by the Gates Foundation (INV-048412 to TJCA) and the MERCK Schistosomiasis Research Grant Initiative (To EN). The authors would also thank Moses Adriko, Fiona Allan, Fred Besigye, Anouk Gouvras, Helmut Haas, Narcis Kabatereine, Sébastien Lambert, Muriel Rabone, Claire Stanley and Edridah Tukahebwa for their prior contributions that helped seed this initiative.

