## Supplementary Files for "TRPtracker: a community database for monitoring praziquantel sensitivity at TRPM_PZQ_ variants"

### Supplementary Material

**Supplementary Table 1.** Functional profiling data for ~170 *Schistosoma mansoni* $TRPM_{PZQ}$ lab mutants and field variants profile using a $Ca^{2+}$ reporter assay. The potency ($EC_{50}$) and efficacy ($B_{max}$) of (±)-PZQ at each $TRPM_{PZQ}$ variant is described together with standard deviation. Relative activity (RA) is a measure of the sensitivity of each $TRPM_{PZQ}$ variant to PZQ, represented in terms of the following categories: 'wild type' (RA, 0.6 – 2, green), 'sensitized' (RA, >2, blue), 'decreased PZQ sensitivity' (RA >0 but <0.6, orange) and complete loss of sensitivity, 'null' (RA = 0, red). Additional information about the natural variants can be found in the associated references, indexed as 1 = Ref. [35], 2 = Ref. [37], 3 = Ref. [36].

**Supplementary Figure 1.** Conservation of residues within a region of the MHR1/2 region that is well conserved between $TRPM_{PZQ}$ orthologs.

**Supplementary Figure 2.** Conservation of phenylalanine residue in the S4 linker (F1521 in *Sm*.$TRPM_{PZQ}$) and tryptophan residue in the TRP domain (W1667A in *Sm*.$TRPM_{PZQ}$) across various $TRPM_{PZQ}$ orthologs.

| Residue | Type | Mutation | Domain | $EC_{50}$ (μM) | *error* | $B_{max}$ (%) | *error* | RA | Effect on $TRPM_{PZQ}$ | Ref |
|---|---|---|---|---|---|---|---|---|---|---|
| wild type | - | - | | **0.74** | | **100** | | **1.000** | Wild type | |
| 102 | Natural Variant | L102V | MHR1/2 | 0.49 | *0.13* | 112.3 | *6.9* | 1.705 | Wild type | 1 |
| 105 | Natural Variant | T105N | MHR1/2 | 0.78 | *0.02* | 84.5 | *4.6* | 0.800 | Wild type | 1 |
| 107 | Lab Mutant | F107A | MHR1/2 | n/a | | 0.0 | | 0.000 | No Response | |
| 108 | Lab Mutant | G108A | MHR1/2 | n/a | | 0.0 | | 0.000 | No Response | |
| 109 | Lab Mutant | T109A | MHR1/2 | 0.76 | *0.13* | 125.9 | *8.2* | 1.224 | Wild type | |
| 110 | Lab Mutant | D110A | MHR1/2 | 0.89 | *0.21* | 127.9 | *5.6* | 1.062 | Wild type | |
| 111 | Lab Mutant | F111A | MHR1/2 | 1.40 | *0.16* | 41.1 | *2.5* | 0.217 | Decrease in Sensitivity | |
| 112 | Lab Mutant | E112A | MHR1/2 | 0.28 | *0.04* | 115.2 | *3.1* | 3.040 | Sensitizes | |
| 145 | Lab Mutant | M145A | MHR1/2 | 0.58 | *0.10* | 67.1 | *8.8* | 0.855 | Wild type | |
| 216 | Lab Mutant | G216A | MHR1/2 | 1.21 | *0.71* | 89.3 | *19.3* | 0.545 | Decrease in Sensitivity | |
| 216 | Lab Mutant | G216Y | MHR1/2 | 0.52 | *0.17* | 72.6 | *5.5* | 1.031 | Wild type | |
| 218 | Lab Mutant | I218A | MHR1/2 | 0.53 | *0.10* | 81.1 | *4.2* | 1.131 | Wild type | |
| 219 | Lab Mutant | K219A | MHR1/2 | 1.09 | *0.30* | 98.1 | *14.0* | 0.665 | Wild type | |
| 222 | Lab Mutant | R222A | MHR1/2 | 0.59 | *0.01* | 76.0 | *8.4* | 0.952 | Wild type | |
| 224 | Lab Mutant | D224A | MHR1/2 | 0.78 | *0.12* | 110.4 | *2.8* | 1.046 | Wild type | |
| 226 | Lab Mutant | E226A | MHR1/2 | 0.84 | *0.23* | 95.3 | *6.2* | 0.838 | Wild type | |
| 228 | Lab Mutant | R228A | MHR1/2 | 0.62 | *0.10* | 105.3 | *7.6* | 1.255 | Wild type | |
| 244 | Lab Mutant | T244A | MHR1/2 | 0.76 | *0.14* | 118.0 | *9.1* | 1.147 | Wild type | |
| 245 | Lab Mutant | T245A | MHR1/2 | 0.90 | *0.17* | 179.7 | *20.1* | 1.475 | Wild type | |
| 246 | Lab Mutant | D246A | MHR1/2 | 0.53 | *0.21* | 78.0 | *6.0* | 1.087 | Wild type | |
| 252 | Lab Mutant | E252A | MHR1/2 | 0.75 | *0.20* | 135.8 | *6.0* | 1.346 | Wild type | |
| 254 | Lab Mutant | E254A | MHR1/2 | 0.73 | *0.10* | 81.0 | *3.1* | 0.820 | Wild type | |
| 258 | Lab Mutant | R258A | MHR1/2 | 0.62 | *0.13* | 138.0 | *10.2* | 1.644 | Wild type | |
| 372 | Natural Variant | A372T | MHR1/2 | 0.47 | *0.07* | 106.4 | *2.8* | 1.673 | Wild type | 2 |
| 374 | Natural Variant | V374I | MHR1/2 | 0.47 | *0.08* | 98.2 | *2.7* | 1.558 | Wild type | 2 |
| 713 | Natural Variant | S713F | MHR3/4 | 0.66 | *0.07* | 83.0 | *2.0* | 0.929 | Wild type | 1 |
| 902 | Natural Variant | E902G | MHR3/4 | 0.48 | *0.06* | 101.8 | *2.3* | 1.584 | Wild type | 2 |
| 915 | Natural Variant | S915C | MHR3/4 | 0.45 | *0.07* | 89.0 | *2.2* | 1.455 | Wild type | 2 |
| 919 | Natural Variant | K919T | MHR3/4 | 0.56 | *0.09* | 85.4 | *2.7* | 1.127 | Wild type | 1 |
| 989 | Natural Variant | P989S | MHR3/4 | 0.41 | *0.09* | 116.0 | *3.7* | 2.075 | Sensitizes | 3 |
| 992 | Natural Variant | F992L | MHR3/4 | 0.49 | *0.10* | 74.9 | *1.0* | 1.129 | Wild type | 1 |
| 993 | Lab Mutant | R993A | MHR3/4 | 0.25 | *0.07* | 100.5 | *1.6* | 2.969 | Sensitizes | |
| 1005 | Natural Variant | T1005I | MHR3/4 | 0.52 | *0.10* | 117.8 | *31.0* | 1.674 | Wild type | 1 |
| 1054 | Natural Variant | F1054C | MHR3/4 | 0.45 | *0.08* | 117.7 | *3.6* | 1.920 | Wild type | 3 |
| 1057 | Natural Variant | E1057N | MHR3/4 | 0.55 | *0.07* | 101.4 | *2.4* | 1.364 | Wild type | 2 |
| 1068 | Natural Variant | M1068I | MHR3/4 | 0.59 | *0.11* | 94.7 | *3.5* | 1.186 | Wild type | 1 |
| 1205 | Lab Mutant | K1205Q | MHR3/4 | 0.69 | *0.10* | 73.2 | *11.5* | 0.784 | Wild type | |
| 1276 | Lab Mutant | Q1276I | MHR3/4 | 0.07 | *0.01* | 77.0 | *2.7* | 7.791 | Sensitizes | |
| 1276 | Lab Mutant | Q1276N | MHR3/4 | 0.18 | *0.01* | 73.7 | *2.3* | 3.025 | Sensitizes | |
| 1280 | Lab Mutant | T1280V | | 0.32 | *0.03* | 176.7 | *2.8* | 4.079 | Sensitizes | |
| 1283 | Lab Mutant | W1283A | | n/a | | 0.0 | | 0.000 | No Response | |
| 1290 | Lab Mutant | K1290Q | | 0.31 | *0.03* | 131.6 | *8.8* | 3.136 | Sensitizes | |
| 1295 | Lab Mutant | R1295Q | | 0.33 | *0.04* | 106.8 | *5.5* | 2.427 | Sensitizes | |
| 1321 | Natural Variant | N1321S | | 0.58 | *0.05* | 91.8 | *1.8* | 1.169 | Wild type | 2 |
| 1341 | Natural Variant | S1341N | | 0.80 | *0.04* | 111.2 | *2.6* | 1.027 | Wild type | 1 |
| 1346 | Lab Mutant | K1346A | | 0.83 | *0.10* | 64.5 | *0.9* | 0.574 | Decrease in Sensitivity | |
| 1347 | Lab Mutant | Y1347A | | 0.97 | *0.09* | 108.2 | *14.1* | 0.824 | Wild type | |
| 1349 | Lab Mutant | L1349A | | 1.02 | *0.23* | 100.0 | *10.0* | 0.724 | Wild type | |
| 1375 | Natural Variant | R1375H | | 0.86 | *0.06* | 66.5 | *1.6* | 0.571 | Decrease in Sensitivity | 2 |
| 1384 | Lab Mutant | R1384A | VSLD | n/a | | 0.0 | | 0.000 | No Response | |
| 1385 | Lab Mutant | F1385A | VSLD | 0.72 | *0.09* | 85.6 | *10.3* | 0.878 | Wild type | |
| 1388 | Lab Mutant | N1388H | VSLD | 0.67 | *0.07* | 93.6 | *2.1* | 1.032 | Wild type | |
| 1388 | Lab Mutant | N1388Q | VSLD | 1.20 | *0.50* | 48.7 | *0.6* | 0.300 | Decrease in Sensitivity | |
| 1388 | Lab Mutant | N1388C | VSLD | 21.60 | *4.10* | 29.1 | *3.0* | 0.010 | Decrease in Sensitivity | |
| 1388 | Lab Mutant | N1388S | VSLD | 24.50 | *5.00* | 29.3 | *1.3* | 0.009 | Decrease in Sensitivity | |
| 1388 | Lab Mutant | N1388A | VSLD | n/a | | 0.0 | | 0.000 | No Response | |
| 1388 | Lab Mutant | N1388T | VSLD | n/a | | 0.0 | | 0.000 | No Response | |
| 1389 | Lab Mutant | T1389V | VSLD | 0.27 | *0.05* | 132.8 | *1.4* | 3.634 | Sensitizes | |

| | | | | | | | | | | |
|---|---|---|---|---|---|---|---|---|---|---|
| 1389 | Lab Mutant | T1389I | VSLD | 0.63 | 0.10 | 41.8 | 2.8 | 0.490 | Decrease in Sensitivity | |
| 1389 | Lab Mutant | T1389S | VSLD | 1.08 | 0.20 | 81.2 | 8.0 | 0.555 | Decrease in Sensitivity | |
| 1389 | Lab Mutant | T1389A | VSLD | 1.85 | 0.84 | 46.6 | 3.1 | 0.186 | Decrease in Sensitivity | |
| 1391 | Lab Mutant | S1391A | VSLD | 0.83 | 0.28 | 75.1 | 4.2 | 0.668 | Wild type | |
| 1392 | Lab Mutant | Y1392A | VSLD | n/a | | 0.0 | | 0.000 | No Response | |
| 1399 | Natural Variant | F1399Y | VSLD | 0.85 | 0.29 | 114.0 | 3.2 | 0.986 | Wild type | 1 |
| 1405 | Natural Variant | V1405I | VSLD | 0.56 | 0.02 | 109.4 | 16.5 | 1.449 | Wild type | 1 |
| 1416 | Natural Variant | Y1416F | VSLD | 0.47 | 0.08 | 108.2 | 3.0 | 1.694 | Wild type | 3 |
| 1420 | Lab Mutant | A1420L | VSLD | 0.89 | 0.40 | 135.3 | 4.7 | 1.123 | Wild type | |
| 1421 | Lab Mutant | W1421A | VSLD | n/a | | 0.0 | | 0.000 | No Response | |
| 1424 | Lab Mutant | T1424S | VSLD | 0.24 | 0.05 | 134.2 | 2.9 | 4.130 | Sensitizes | |
| 1424 | Lab Mutant | T1424A | VSLD | 1.39 | 0.29 | 82.1 | 5.3 | 0.436 | Decrease in Sensitivity | |
| 1425 | Lab Mutant | L1425A | VSLD | 2.41 | 0.41 | 61.9 | 9.0 | 0.190 | Decrease in Sensitivity | |
| 1425 | Lab Mutant | L1425Y | VSLD | 6.70 | 1.30 | 73.6 | 2.1 | 0.081 | Decrease in Sensitivity | |
| 1428 | Lab Mutant | E1428A | VSLD | n/a | | 0.0 | | 0.000 | No Response | |
| 1429 | Lab Mutant | E1429A | VSLD | 33.10 | 3.50 | 11.9 | 2.0 | 0.003 | Decrease in Sensitivity | |
| 1431 | Lab Mutant | K1431A | VSLD | 1.78 | 0.30 | 77.2 | 3.4 | 0.320 | Decrease in Sensitivity | |
| 1432 | Lab Mutant | Q1432A | VSLD | 1.03 | 0.37 | 70.4 | 4.0 | 0.505 | Decrease in Sensitivity | |
| 1435 | Lab Mutant | W1435A | VSLD | 0.90 | 0.20 | 155.1 | 5.1 | 1.273 | Wild type | |
| 1436 | Lab Mutant | A1436L | VSLD | 2.10 | 0.13 | 41.8 | 6.8 | 0.147 | Decrease in Sensitivity | |
| 1445 | Lab Mutant | T1445A | VSLD | n/a | | 0.0 | | 0.000 | No Response | |
| 1446 | Lab Mutant | Y1446A | VSLD | n/a | | 0.0 | | 0.000 | No Response | |
| 1451 | Lab Mutant | W1451A | VSLD | n/a | | 0.0 | | 0.000 | No Response | |
| 1452 | Lab Mutant | N1452A | VSLD | 2.50 | 0.90 | 41.9 | 2.8 | 0.124 | Decrease in Sensitivity | |
| 1455 | Lab Mutant | D1455A | VSLD | n/a | | | | | No Response | |
| 1458 | Lab Mutant | G1458A | VSLD | 1.20 | 0.40 | 98.1 | 9.4 | 0.604 | Wild type | |
| 1459 | Lab Mutant | L1459A | VSLD | n/a | | 0.0 | | 0.000 | No Response | |
| 1476 | Natural Variant | L1476I | VSLD | 0.96 | 0.38 | 103.0 | 9.8 | 0.796 | Wild type | 1 |
| 1511 | Lab Mutant | F1511A | VSLD | n/a | | 0.0 | | 0.000 | No Response | |
| 1514 | Lab Mutant | R1514A | VSLD | n/a | | 0.0 | | 0.000 | No Response | |
| 1517 | Lab Mutant | Y1517A | VSLD | n/a | | 0.0 | | 0.000 | No Response | |
| 1517 | Lab Mutant | Y1517H | VSLD | n/a | | 0.0 | | 0.000 | No Response | |
| 1517 | Lab Mutant | Y1517E | VSLD | n/a | | 0.0 | | 0.000 | Decrease in Sensitivity | |
| 1518 | Lab Mutant | T1518I | VSLD | 0.17 | 0.07 | 147.5 | 2.8 | 6.410 | Sensitizes | |
| 1518 | Lab Mutant | T1518S | VSLD | 0.55 | 0.04 | 116.4 | 1.0 | 1.563 | Wild type | |
| 1518 | Lab Mutant | T1518A | VSLD | 0.53 | 0.20 | 121.8 | 11.1 | 1.698 | Wild type | |
| 1518 | Lab Mutant | T1518L | VSLD | 0.92 | 0.21 | 135.9 | 4.4 | 1.091 | Wild type | |
| 1520 | Lab Mutant | S1520A | VSLD | 2.90 | 0.60 | 51.6 | 2.1 | 0.131 | Decrease in Sensitivity | |
| 1521 | Lab Mutant | F1521A | pore/TRP | 0.09 | 0.00 | 123.0 | 2.3 | 10.530 | Sensitizes | |
| 1522 | Lab Mutant | H1522Q | pore/TRP | 27.50 | 25.20 | 27.0 | 4.8 | 0.007 | Decrease in Sensitivity | |
| 1523 | Lab Mutant | I1523A | pore/TRP | n/a | | 0.0 | | 0.000 | No Response | |
| 1525 | Lab Mutant | L1525A | pore/TRP | n/a | | 0.0 | | 0.000 | No Response | |
| 1526 | Lab Mutant | G1526C | pore/TRP | n/a | | 0.0 | | 0.000 | No Response | |
| 1526 | Lab Mutant | G1526S | pore/TRP | n/a | | 0.0 | | 0.000 | No Response | |
| 1527 | Lab Mutant | P1527A | pore/TRP | n/a | | 0.0 | | 0.000 | No Response | |
| 1543 | Lab Mutant | F1543L | pore/TRP | 0.97 | 0.10 | 17.0 | 2.1 | 0.129 | Decrease in Sensitivity | |
| 1551 | Lab Mutant | M1551I | pore/TRP | 0.70 | 0.10 | 38.2 | 5.4 | 0.404 | Decrease in Sensitivity | |
| 1554 | Natural Variant | Y1554C | pore/TRP | n/a | | | | 0.000 | No Response | 1 |
| 1600 | Lab Mutant | F1600A | pore/TRP | n/a | | 0.0 | | 0.000 | No Response | |
| 1601 | Lab Mutant | G1601A | pore/TRP | 0.65 | 0.10 | 117.6 | 0.7 | 1.337 | Wild type | |
| 1602 | Lab Mutant | D1602E | pore/TRP | 0.46 | 0.10 | 104.1 | 2.4 | 1.672 | Wild type | |
| 1602 | Lab Mutant | D1602A | pore/TRP | 9.17 | 3.60 | 36.5 | 1.1 | 0.029 | Decrease in Sensitivity | |
| 1606 | Lab Mutant | D1606A | pore/TRP | 2.80 | 1.50 | 66.6 | 2.5 | 0.176 | Decrease in Sensitivity | |
| 1609 | Lab Mutant | Q1609A | pore/TRP | 0.76 | 0.20 | 80.6 | 10.7 | 0.784 | Wild type | |
| 1615 | Lab Mutant | C1615A | pore/TRP | n/a | | 0.0 | | 0.000 | No Response | |
| 1620 | Lab Mutant | C1620A | pore/TRP | 0.83 | 0.13 | 38.1 | 1.2 | 0.339 | Decrease in Sensitivity | |
| 1624 | Natural Variant | T1624K | pore/TRP | 1.41 | 0.23 | 79.6 | 5.5 | 0.417 | Decrease in Sensitivity | 1 |
| 1653 | Lab Mutant | S1653A | pore/TRP | 0.41 | 0.02 | 128.2 | 2.8 | 2.339 | Sensitizes | |
| 1656 | Lab Mutant | Y1656F | pore/TRP | 0.40 | 0.10 | 192.0 | 4.1 | 3.546 | Sensitizes | |
| 1657 | Lab Mutant | E1657A | pore/TRP | n/a | | 0.0 | | 0.000 | No Response | |

| | | | | | | | | | | |
|---|---|---|---|---|---|---|---|---|---|---|
| 1663 | Lab Mutant | S1663V | pore/TRP | n/a | | 0.0 | | 0.000 | No Response | |
| 1667 | Lab Mutant | W1667A | pore/TRP | 0.06 | *0.01* | 100.0 | *1.6* | 12.849 | Sensitizes | |
| 1668 | Lab Mutant | N1668A | pore/TRP | 0.64 | *0.20* | 101.1 | *2.7* | 1.167 | Wild type | |
| 1669 | Lab Mutant | Y1669A | pore/TRP | 0.73 | *0.10* | 64.5 | *5.7* | 0.653 | Wild type | |
| 1670 | Lab Mutant | Q1670A | pore/TRP | n/a | | 0.0 | | 0.000 | No Response | |
| 1670 | Natural Variant | Q1670K | pore/TRP | n/a | | 0.0 | | 0.000 | No Response | 1 |
| 1671 | Lab Mutant | R1671K | pore/TRP | 0.26 | *0.05* | 167.9 | *6.2* | 4.770 | Sensitizes | |
| 1671 | Lab Mutant | R1671E | pore/TRP | n/a | | 0.0 | | 0.000 | No Response | |
| 1671 | Lab Mutant | R1671H | pore/TRP | 0.37 | *0.03* | 47.6 | *1.0* | 0.950 | Wild type | |
| 1671 | Lab Mutant | R1671Q | pore/TRP | 2.20 | *0.40* | 16.5 | *1.8* | 0.055 | Decrease in Sensitivity | |
| 1671 | Lab Mutant | R1671A | pore/TRP | n/a | | 0.0 | | 0.000 | No Response | |
| 1672 | Lab Mutant | Y1672A | pore/TRP | 1.00 | *0.70* | 42.2 | *5.7* | 0.312 | Decrease in Sensitivity | |
| 1673 | Lab Mutant | Q1673K | pore/TRP | 0.65 | *0.06* | 55.3 | *1.2* | 0.628 | Wild type | |
| 1674 | Lab Mutant | M1674E | pore/TRP | n/a | | 0.0 | | 0.000 | No Response | |
| 1674 | Lab Mutant | M1674R | pore/TRP | n/a | | 0.0 | | 0.000 | No Response | |
| 1674 | Lab Mutant | M1674A | pore/TRP | n/a | | 0.0 | | 0.000 | No Response | |
| 1676 | Lab Mutant | N1676A | pore/TRP | 0.46 | *0.10* | 121.1 | *2.6* | 1.945 | Wild type | |
| 1677 | Lab Mutant | D1677Y | pore/TRP | 0.18 | *0.04* | 159.5 | *8.4* | 6.547 | Sensitizes | |
| 1677 | Lab Mutant | D1677L | pore/TRP | n/a | | 0.0 | | 0.000 | No Response | |
| 1677 | Lab Mutant | D1677A | pore/TRP | 1.40 | *0.30* | 27.0 | *1.0* | 0.142 | Decrease in Sensitivity | |
| 1677 | Lab Mutant | D1677Q | pore/TRP | 4.78 | *0.77* | 16.9 | *2.8* | 0.026 | Decrease in Sensitivity | |
| 1677 | Lab Mutant | D1677E | pore/TRP | n/a | | 0.0 | | 0.000 | No Response | |
| 1677 | Lab Mutant | D1677N | pore/TRP | n/a | | 0.0 | | 0.000 | No Response | |
| 1678 | Lab Mutant | Y1678F | pore/TRP | 4.18 | *1.30* | 67.1 | *2.5* | 0.119 | Decrease in Sensitivity | |
| 1678 | Lab Mutant | Y1678A | pore/TRP | n/a | | 0.0 | | 0.000 | No Response | |
| 1680 | Lab Mutant | H1680Y | pore/TRP | 0.87 | *0.11* | 112.0 | *5.8* | 0.951 | Wild type | |
| 1680 | Lab Mutant | H1680W | pore/TRP | 1.40 | *0.20* | 95.8 | *5.4* | 0.506 | Decrease in Sensitivity | |
| 1680 | Lab Mutant | H1680A | pore/TRP | 1.60 | *0.30* | 96.6 | *5.4* | 0.446 | Decrease in Sensitivity | |
| 1681 | Lab Mutant | R1681A | pore/TRP | 2.10 | *0.70* | 63.0 | *7.1* | 0.222 | Decrease in Sensitivity | |
| 1681 | Lab Mutant | R1681K | pore/TRP | 0.99 | *0.10* | 107.5 | *2.1* | 0.802 | Wild type | |
| 1682 | Lab Mutant | S1682A | pore/TRP | 0.55 | *0.20* | 85.6 | *5.1* | 1.150 | Wild type | |
| 1686 | Lab Mutant | P1686A | pore/TRP | 6.70 | *2.20* | 18.2 | *1.1* | 0.020 | Decrease in Sensitivity | |
| 1687 | Lab Mutant | P1687A | pore/TRP | n/a | | 0.0 | | 0.000 | No Response | |
| 1688 | Lab Mutant | I1688A | | 0.80 | *0.20* | 129.8 | *20.9* | 1.199 | Wild type | |
| 1690 | Lab Mutant | I1690A | | 1.10 | *0.10* | 83.7 | *7.1* | 0.562 | Decrease in Sensitivity | |
| 1692 | Lab Mutant | W1692A | | 0.76 | *0.09* | 87.6 | *4.7* | 0.852 | Wild type | |
| 1693 | Lab Mutant | H1693A | | n/a | | 0.0 | | 0.000 | No Response | |
| 1696 | Lab Mutant | E1696A | | 1.20 | *0.24* | 85.3 | *6.6* | 0.525 | Decrease in Sensitivity | |
| 1703 | Lab Mutant | N1703A | | 1.00 | *0.20* | 129.2 | *10.9* | 0.955 | Wild type | |
| 1704 | Lab Mutant | Q1704A | | 1.50 | *0.50* | 114.1 | *6.6* | 0.562 | Decrease in Sensitivity | |
| 1705 | Lab Mutant | C1705A | | 1.50 | *0.40* | 79.0 | *6.5* | 0.389 | Decrease in Sensitivity | |
| 1840 | Natural Variant | R1840L | | 0.37 | *0.08* | 112.1 | *3.1* | 2.239 | Sensitizes | 3 |
| 1843 | Natural Variant | R1843Q | | 1.14 | *0.30* | 90.8 | *9.1* | 0.589 | Decrease in Sensitivity | 1 |
| 1870 | Natural Variant | Q1870H | | 0.71 | *0.09* | 102.1 | *3.5* | 1.062 | Wild type | 2 |
| 2170 | Natural Variant | E2170D | nudix | 0.68 | *0.05* | 86.6 | *1.8* | 0.941 | Wild type | 2 |
| 2195 | Lab Mutant | D2195A | nudix | 0.60 | *0.20* | 83.5 | *8.2* | 1.028 | Wild type | |
| 2196 | Lab Mutant | Q2196A | nudix | 1.04 | *0.30* | 83.3 | *23.9* | 0.592 | Decrease in Sensitivity | |
| 2197 | Lab Mutant | L2197A | nudix | 1.14 | *0.40* | 96.9 | *11.2* | 0.628 | Wild type | |
| 2200 | Lab Mutant | D2200A | nudix | 0.90 | *0.15* | 104.7 | *20.9* | 0.859 | Wild type | |
| 2220 | Natural Variant | Q2220H | nudix | 0.49 | *0.06* | 86.8 | *1.9* | 1.309 | Wild type | 1 |
| 2255 | Lab Mutant | Y2255A | nudix | 0.72 | *0.10* | 129.4 | *3.5* | 1.328 | Wild type | |

```
Trematodes     S.mansoni             NSETPDSILRDILKRKWGLKPPTLIITVFGTDFEKKRKLKMIFKKGLWKAAES-GCWIVT
               S.bovis               NSDTPDSVLRDILKRKWGLKPPTLIITVFGTDFEKKRKLKMIFKKGLWKAAES-GCWIVT
               S.curassoni           NSDTPDSVLRDILKRKWGLKPPTLIITVFGTDFEKKRKLKMIFKKGLWKAAES-GCWIVT
               S.haematobium         NSDTPDSVLRDILKRKWGLKPPTLIITVFGTDFEKKRKLKMIFKKGLWKAAES-GCWIVT
               S.japonicum           NSDTADSVLRDILKRKWGLKPPTLIITVFGTDFEKKRKLKMIFKKGLWKAAES-GCWIVT
               S.margrebowiei        NSDTPDSVLRDILKRKWGLKPPTLIITVFGTDFEKKRKLKMIFKKGLWKAAES-GCWIVT
               S.rodhaini            NSDTPDSILRDILKRKWGLKPPTLIITVFGTDFEKKRKLKMIFKKGLWKAAES-GCWIVT
               T.regenti             NSDTADSVLRDILKRKWGLKPPTLIITVFGTDFEKKRKLKMIFKKGLWKAAES-GCWIVT
               E.caproni             DGSTTDQVIRDLLKRKWGLKPPTLIITVFGTDFEKKRKLKMIFKKGLWKAAES-GCWIVT
               F.gigantica           DSNTNDQVIRDLLKRKWGLKPPTLIITVFGTDFEKKRKLKMIFKKGLWKAAES-GCWIVT
               F.hepatica            DSSTNDQVIRDLLKRKWGLKPPTLIITVFGTDFEKKRKLKMIFKKGLWKAAES-GCWIVT
               F.buski               DSNTNDYVIRDLLKRKWGLKPPTLIITVFGTDFEKKRKLKMIFKKGLWKAAES-GCWIVT
               A.winterbourni        ETTTNDSVIRDLLKRKWGLKPPTLIITVFGTDFEKKRKLKMIFKKGLWKAAES-GCWIVT
               P.heterotremus        DTATHDTVIRDLLKRKWGLKPPTLIITVFGTDFEKKRKLKMIFKKGLWKAAES-GCWIVT
               C.sinensis            DSSTNDMVIRDLLKRKWGLKPPTLIITVFGTDFEKKRKLKMIFKKGLWKAAES-GCWIVT
               O.viverrine           DSSTNDTVIRDLLKRKWGLKPPTLIITVFGTDFEKKRKLKMIFKKGLWKAAES-GCWIVT
Monogeneans    G.bullartarudis       PENTPDSTIRDLLKRKWFLKPPTLIITVFGTNFEKKRKLKMIFKKGLWKAAES-GCWIVT
               G.salaris             PDNTPDSTIRDLLKRKWSLKPPTLIITVFGTNFEKKRKLKMIFKKGLWKAAES-GCWIVT
Cestodes       E.canadensis          SSNMSDSAIRDLLQRRWGLKPPTLIITVFGADFEKKRKLKMIFKKGLWKAADS-GCWIVT
               E.granulosus          SSSMSDSAIRDLLQRRWSLKPPTLIITVFGTDFEKKRKLKMIFKKGLWKAADS-GCWIVT
               E.multilocularis      SSSMSDSAIRDLLQRRWGLKPPTLIITVFGTDFEKKRKLKMIFKKGLWKAADS-GCWIVT
               H.taeniaeformis       SPCISDSAIRDLLQRRWGLKPPTLIITVYGTDFEKKRKLKMIFKKGLWKAADS-GCWIVT
               T.asiatica            SPCMSDSAIRDLLQRRWGLKPPTLIITVYGTDFEKKRKLKMIFKKGLWKAADS-GCWIVT
               T.multiceps           SPFMSDSAIRDLLQRRWGLKPPTLIITVYGTDFEKKRKLKMIFKKGLWKAADS-GCWIVT
               T.saginata            SPCMSDSAIRDLLQRRWGLKPPTLIITVYGTDFEKKRKLKMIFKKGLWKAADS-GCWIVT
               T.solium              SPCMSDSAIRDLLQRRWGLKPPTLIITVYGTDFEKKRKLKMIFKKGLWKAADS-GCWIVT
               H.diminuta            SPYMSDSAIRDLLQRRWGLKPPTLIITVFGTDFEKKRKLKMIFKKGLWKAADS-GCWIVT
               H.microstoma          SPHMSDNAIRDLLQRRWGLRPPTLIITVFGTDFEKKRKLKMIFKKGLWKAADS-GCWIVT
               R.nana                SPHMSDSAIRDLLQRRWGLRPPTLIITVFGTDFEKKRKLKMIFKKGLWKAADS-GCWIVT
               M.corti               SPYMTDSAIRDLLQRRWGLKPPTLIITVFGTDFEKKRKLKMIFKKGLWKAADS-GCWIVT
               M.expansa             SPHMSDSAIRDLLQRRWGLKPPTLIITVYGTDFEKKRKLKMIFKKGLWKAADS-GCWIVT
               S.solidus             SNSVLDSAIRDLLQRRWALKPPTLIITVYGTDFEKKRKLKMIFKKGLWKAADS-GCWIVT
               S.erinaceieuropaei    SNSVLDSAIRDLLQRRWALKPPTLIITVYGTDFEKKRKLKMIFKKGLWKAADS-GCWIVT
Free Living    M.lignano             DSNTPEEALHELLVKKWEMSRPTLVINIFGGDFEKKRQLKMIFKKGLWKAAESAGCWIVT
                                          :  ::::* ::* :  ***:*.::* :*****:*************:* ******
```

Supplementary Figure 1

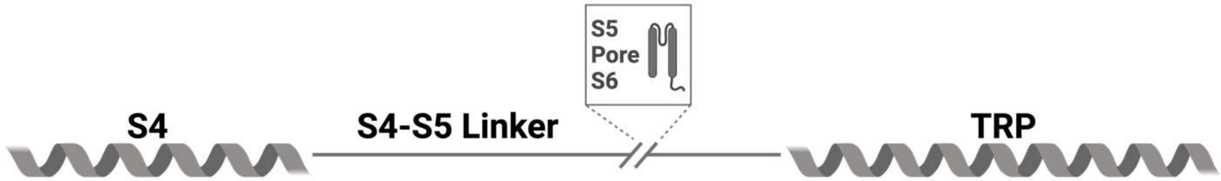


```
S.mansoni             SRICLAISLFTFYIRLMYTFSFHIALGPKLIMIGKMVTNDL…KTYEEIESASTYYWNYQRYQMINDYVHRSPLAPPIII
S.haematobium         SRICLAISLFTFYIRLMYTFSFHIALGPKLIMIGKMVTNDL…KTYEEIESASTYYWNYQRYQMINDYVHRSPLAPPIII
S.japonicum           SRICLAISLFTFYIRLMYTFSFHIALGPKLIMIGKMVTNDL…KTYEEIESASTYYWNYQRYQMINDYVHRSPLAPPIII
E.caproni             SRTCLAISLFVFYTRLMYTFSFHIALGPKLIMIGKMVTNDL…KTYEEIESASTYYWNYQRYQMINDYVHRSPLVPPIII
F.hepatica            SRTCLAISLFVFYTRLMYTFSFHIALGPKLIMIGKMVTNDL…KTYEEIESASTYYWNYQRYQMINDYVHRSPLVPPIII
C.sinensis            SRISLAISLFVFYIRLMYTFSFNIALGPKLIMIGKMVTNDL…KTYEEIESASTYYWNYQRYQMINDYVHRSPLVPPIII
O.viverrine           SRISLAISLFVFYIRLMYTFSFNIALGPKLIMIGKMVTNDL…KTYEEIESASTYYWNYQRYQMINDYVHRSPLVPPIII
G.salaris             SRICLAISLFTFYIRLMYTFSFHIALGPKLIMIGKMVTNDL…KTYEEIESASTYYWNYQRYQMIAEYVHRSPLAPPLIL
E.granulosus          SRICMAFSLFVFYIRLMYSFSFHIALGPKLIMIGKMVTNDL…KTYEEIESASTYYWNYQRYQMIADYVRRSPLVPPLII
E.multilocularis      SRICMAFSLFVFYIRLMYSFSFHIALGPKLIMIGKMVTNDL…KTYEEIESASTYYWNYQRYQMIADYVRRSPLVPPLII
T.multiceps           SRICMAFSLFVFYIRLMYSFSFHIALGPKLIMIGKMVTNDL…KTYEEIESASTYYWNYQRYQMIADYVRRSPLVPPLII
T.solium              SRICMAFSLFVFYIRLMYSFSFHIALGPKLIMIGKMVTNDL…KTYEEIESASTYYWNYQRYQMIADYVRRSPLVPPLII
H.diminuta            SRICMAISLFVFYIRLMYSFSFHIALGPKLIMIGKMVTNDL…KTYEEIESASTYYWNYQRYQMIADYVRRSPLVPPLII
M.corti               SRICMAFSLFVFYIRLMYSFSFHIALGPKLIMIGKMVTNDL…KTYEEIESASTYYWNYQRYQMIADYVRRSPLVPPLII
S.erinaceieuropaei    SRICMAFSLFVFYIRLMYTFSFHIALGPKLIMIGKMVTNDL…KTYEEIESASTYYWNYQRYQMIAEYVRRSPLVPPIII
M.lignano             ARICYAFSLFAFCIRLMYIFSFHIALGPKLIMIGKMVINDL…STYESINQFSALHWNYQRYSMIKEYIERSPLAPPLII
                      :*  .  *:***.*   ****  ***:************** *** .***.*:.  *:  :******.**  :*:.****.**:*:
```

Supplementary Figure 2
